# ET-Pfam: Ensemble transfer learning for protein family prediction

**DOI:** 10.1101/2025.08.02.668111

**Authors:** Sofia Escudero, Sofia Duarte, Rosario Vitale, Emilio Fenoy, Leandro Bugnon, Diego H. Milone, Georgina Stegmayer

**Affiliations:** Research Institute for Signals, Systems and Computational Intelligence, sinc(i), FICH-UNL,CONICET, Ciudad Universitaria UNL, 3000, Santa Fe, Argentina

## Abstract

**Motivation:** Due to the rapid growth of sequence generation, which has surpassed the expert curators ability to manually review and annotate them, the computational annotation of proteins remains a significant challenge in bioinformatics nowadays. The Pfam database contains a large collection of proteins that are nowadays annotated with domain families through multiple sequence alignments and profile Hidden Markov models (pHMMs). However, such computational annotation methods have some limitations such as problems for handling large datasets and the fact that multiple sequence alignments are computationally challenging to compute with high accuracy due to the increase in complexity as the number of sequences and lengths grow. Additionally, each HMM is independently obtained for each family missing the opportunity of learning patterns across families, that is from a complete view of all the dataset. As an alternative, some deep learning (DL) models have been recently proposed, nevertheless with simple representations of the inputs and moderate improvements in performance.

**Results:** In this work we present ET-Pfam, a novel approach based on transfer learning and ensembles of multiple DL classifiers to predict functional families in the Pfam database. Several base DL models are first trained using learned representations from a protein large language model, with different hyperparameters to increase diversity. Then, the base models are integrated using classical ensemble strategies and novel voting approaches by learning weights for each model and for each Pfam family. Results demonstrate that the proposed ET-Pfam method can consistently diminish classification error rates compared to individual DL models, boosting prediction performance. Among the novel ensemble strategies presented here, the learned weights by family voting achieved the best performance, with the lowest error rate (7.00%), significantly surpassing the best individual base model error (12.91%) and three competitors of the state-of-the-art on the same Pfam dataset.

**Availability:** Data and source code are available at https://github.com/sinc-lab/ET-Pfam.

## 1 Introduction

In proteomics, the task of protein family prediction involves identifying similarity and evolutionary relationships between proteins, which is necessary for applications such as drug discovery and functional annotation of genomes [13]. Nowadays, one of the most widely used computational approaches for protein family prediction is sequence alignment. The alignment of multiple sequences having the same or closely related function can be used for modeling protein families and also for determining functions of yet unannotated proteins [19]. However, their computational complexity increases exponentially to the number of sequences [23]. As of June 2025 there are 24,736 protein families and 62,646,683 sequences annotated with their corresponding family domain in Pfam v37.4. However, the UniProtKB database that constitutes the complete world of known proteins up-to-date has 253,635,358 sequence entries. This means that it was possible to annotate the Pfam family of just 25% of the known protein universe.

The protein family annotation task is handled at the protein family database Pfam with multiple sequence alignments and pHMMs [8, 2]. The state-of-the-art in protein domain sequence annotation uses pHMMs to detect domains [9, 16]. Pfam is a comprehensive database of protein families and domains where each family entry contains a seed alignment of a representative set of sequences, from which a pHMM is generated [16]. These pHMM are then used to search against UniProtKB proteomes. Sequence regions that meet a family-specific curated threshold are classified with the corresponding family of the pHMM. However, in many families it is impossible to build a pHMM because the examples available are just a few. Note that HMMs are single-domain generative models, meaning that each pHMM learns the underlying probability distribution from sequences of just one specific family without using information from the others, in order to generate samples that closely resemble training data. Thus, such a learning approach does not take advantage of sequences from several families to learn inner patterns that might be shared among them. Moreover, profiles for training are based on manually curated sequences, and pHMM models cannot learn patterns from insufficient examples that have not been properly aligned. Profile HMMs make simplifying independence assumptions when modeling residues in proteins, since they assume that the observed residues are conditionally independent given the hidden state, which may limit their ability to capture complex dependencies among residues. Thus, there is large interest in the community in the development of other methods that can better recognize more complex or distant evolutionary relationships.

As a completely different approach and inspired by the recent success of deep learning (DL) models in the bioinformatics community [14], alignment-free methods for modeling protein families with DL have appeared. DL models have the capability of learning a model from information shared across several families, and can directly predict annotation from unaligned sequences. In fact, they can capture local and non-local patterns in the sequence, and transfer information across protein families. DL models are discriminative models, thus focus on learning features to define a decision boundary that separates classes within data. Instead of modeling data distribution itself like HMMs, DL models aim to learn features with discriminative information in the training data. Thus, discriminative models are very suited for classification tasks since they can effectively capture the differences between classes. Moreover, DL models are trained looking at the complete sets of training patterns at the same time. These models are capable of inferring inner patterns or hidden rules shared across families, which can be very useful when new sequences have to be annotated. Some preliminary DL models were proposed for the Pfam annotation task [19, 3], however relying on very simple input data representations such as one-hot encoding completed with zero padding. ProtENN [3], an ensemble of convolutional neural networks, was designed to use only a cropped segment with an isolated domain as input, that is, it needs the beginning and ending marks of curated Pfam seeds. Both models were designed to classify one domain per sequence at a time, thus the length of the input sequence may end up being the key feature used for prediction.

On the one hand, it has been shown recently [22] that using transfer learning from pre-trained large language models (LLMs) can reduce prediction errors in comparison to standard methods and DL models with one-hot encoding. Analogously to what happened in natural language processing, several protein language models (pLMs) have appeared in the last five years [26, 6, 20] that take advantage of the vast quantity of unannotated protein sequence data available. Protein language models can capture some aspects of the grammar of the language of life as written in protein sequences [25]. As a matter of fact, pLMs have now turned into an important representation alternative for capturing the information encoded in a protein, thus becoming an increasingly powerful means of advancing protein prediction and annotation [12]. For many applications, alignment-free pLM-based predictions now have become significantly more accurate [25]. A pre-trained pLM, given a raw protein sequence, can calculate a dense feature vector (embedding) that encodes its representation, which can serve as the exclusive input into downstream supervised methods greatly simplifying the modeling. Next, a predictive model can solve the downstream task by learning from embeddings associated with specific target labels. Interestingly, pLMs do not require annotations during pre-training, as they can learn from the raw protein sequences. This capability by which LLMs acquire knowledge by self-supervised learning from a large dataset (for example, UniProtKB), and use it into another more specific task that requires annotated data (for example, Pfam classification) is named transfer learning (TL) [21, 24]. In a review [7], several protein sequence representation learning methods were experimentally benchmarked, resulting in Evolutionary Scale Modeling (ESM) [18] being the best method for the tasks evaluated. Recent evaluations of pLMs for annotating proteins confirmed ESM2 as the best choice to boost protein families classification in the Pfam database [22, 4]. On the other hand, the flexibility, adaptability and exceptional performance of ensemble methods together with DL have spread their application in bioinformatics research [11]. In spite of these techniques being developed independently in machine learning, the recent emergence of ensemble deep learning has prompted new applications. With ensembles, instead of just building a single model, multiple and varied base models are combined to achieve better generalization performance. In the field of bioinformatics, ensembles have shown the capability to deal with small sample sizes, high dimensionality, unbalanced class distribution, and noisy and heterogeneous data generated by biological systems [5].

In this work we propose the combination of two strategies to improve the state-of-the art prediction of protein families in Pfam. First, to leverage the success of TL from pLMs trained in an self-supervised way from largescale protein data. Second, to develop an ensemble of several DL models combined with TL from a pLM, ET-Pfam, developing strategies beyond the classical ensemble voting, such as learned weights by model voting and learned weights by family voting, to boost prediction performance.

## 2 ET-Pfam ensemble model

The architecture of ET-Pfam is shown in Figure 1. First, each protein sequence of length *L* in the dataset is passed through a pLM in order to obtain a corresponding embedding of size *E* × *L*, where *E* is the embedding dimension. Then several base models are trained separately, indicated in the figure with different colors, having the same architecture but different hyperparameters. Each base model has a sampler that generates *N* input windows of length *W* residues from the complete sequence, with its corresponding Pfam family classification label. In training, each window is randomly sampled. In testing, the window is slided with discrete steps along the protein. The base model has a ResNet block with a convolutional (conv1D) layer, a bottleneck, and another conv1D, with *F* filters for the first and the last convolutions, and *F*_*b*_ filters in the bottleneck. After that, a max pooling along the window is performed, obtaining a vector of *F* × 1 elements. Then there is a linear fully connected layer, with *K* output classes (the number of Pfam families), which provides the final prediction for each window. Cross-entropy loss and Adam optimizer are used during training. Each base model can be trained with its own learning rate *l*_*r*_, and uses a different window length *W* in the sampler. For the final output, a Pfam family prediction can be provided according to several strategies to ensemble the output scores of the base models.

**Figure 1.**
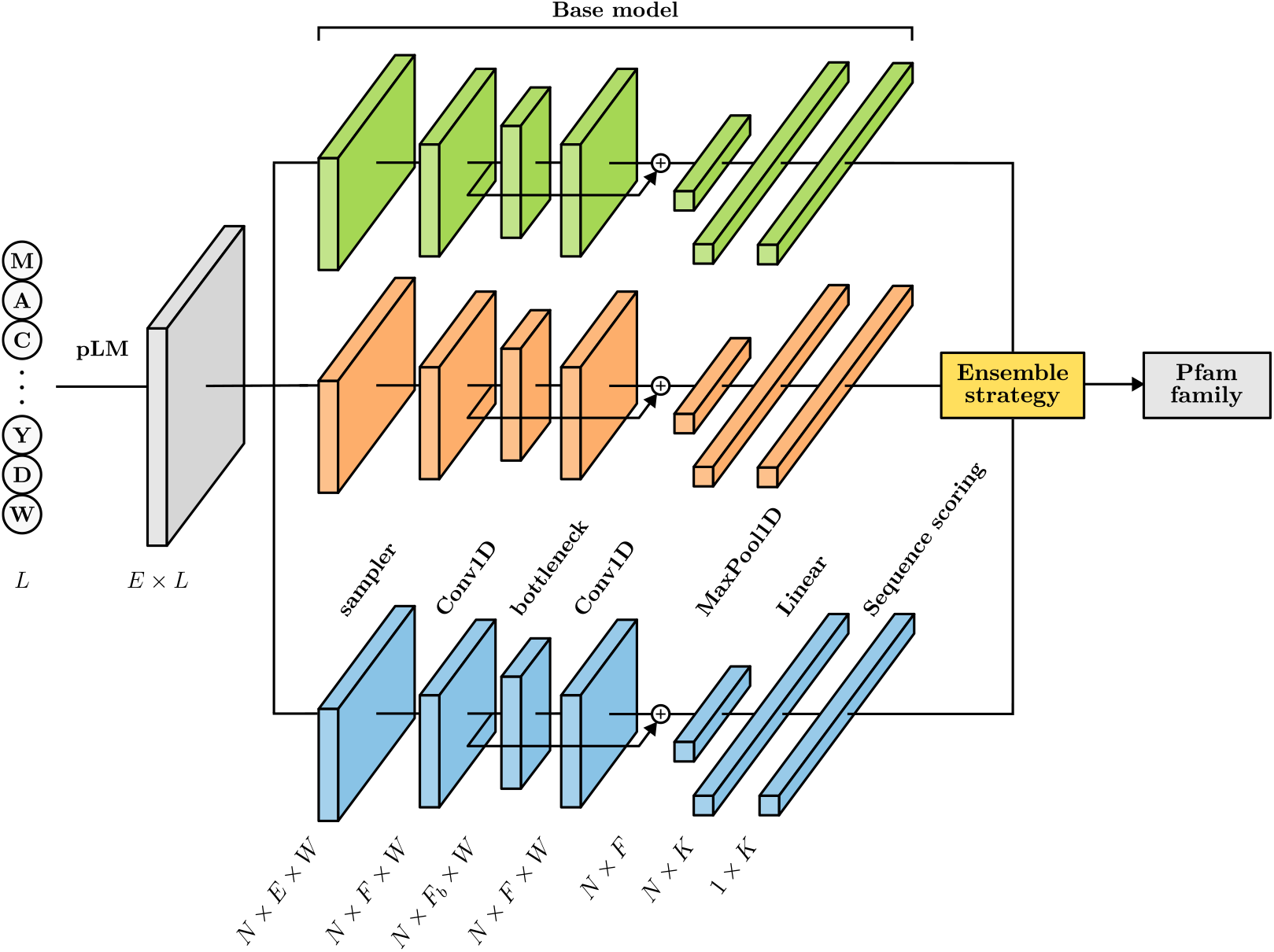
ET-Pfam architecture for Pfam family prediction in full sequences. Each input protein is represented with an embedding obtained from a pLM. The sequence of length *L* is sampled in smaller windows *W*, with its corresponding family class. Several base models, indicated in the figure with different colors, receive the same window embeddings and are trained separately. Each DL base model has the same architecture but different hyperparameters. At the output, using the scores of base models the prediction of a Pfam family *K* can be provided according to different ensemble strategies: simple voting, score voting, learned weights by model voting and learned weights by family voting.

### 2.1 Base models scores

In this proposal is it important to highlight that, unlike other approaches, here the sampler extracts windows of the full protein and the convolutions are made inside each window. Given a window centered in the *n*-th residue or the protein, let be *s*_*ik*_(*n*) the output score of the *i*-th model of the ensemble, for the *k*-th output class (Pfam family). For each test sequence, the simplest way of obtaining the final class score is by considering the highest score of an input window positioned at the central residue *n*_*c*_. Given the beginning (*n*_*b*_), ending (*n*_*e*_) and the central window coordinate 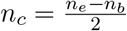, the central window score (CwS) is

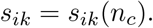

A better prediction can be obtained by considering the scores of the sliding windows along the test sequence. Then, the output score for each class *k* will be the maximum score along all positions *n*. That is, *s*_*ik*_ = max_*n*_ *s*_*ik*_(*n*). However, this value can be just a single peak within the full range of the test sequence. In order to better take into account all the scores within the complete test sequence, we propose two alternatives.

The first alternative is, given the beginning (*n*_*b*_) and ending (*n*_*e*_) residues, to calculate the area under the *s*_*i,k*_(*n*) along *n*, that is the sliding window area (SwA) as

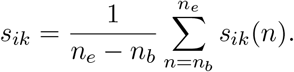

The second alternative is to account for the coverage of the score, considering the times that each class was the winner along all the amino acids of the test sequence. The sliding window coverage (SwC) can be written as

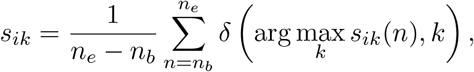

where the Kroneker delta is *δ*(*k*^*∗*^, *k*) = 1 when *k*^*∗*^ = *k* and 0 otherwise.

Finally, for each test sequence, whatever the alternative used to obtain *s*_*ik*_, the predicted family *κ*_*i*_ of the base model *i* can be obtained simply with the maximum score *κ*_*i*_ = arg max_*k*_ *s*_*ik*_.

### 2.2 ET-Pfam ensemble voting strategies

At the output of ET-Pfam, an ensemble prediction of a determined Pfam family *κ*^*∗*^ can be provided according to different ensemble strategies: simple voting, score voting, learned weights by model voting and learned weights by family voting.

#### Simple voting

In this voting strategy, for each possible class *k*, the sum of the base model votes is obtained and then the most voted class is selected. That is, the final predicted family *κ*^*∗*^ is

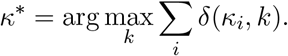

#### Score voting

Instead of considering that each base model votes with 1 or 0, the score of the output class prediction can be used to balance the voting. Therefore, the sum of the output scores of the base models is obtained, and the class with the maximum sum of scores is selected as the output prediction of the ensemble. This can be expressed simply as

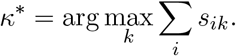

#### Learned weights by model (LWM) voting

Due to the fact that not all base models are equally good at predicting during training, in this schema the score of each base model is weighted according

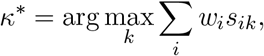

to where the weights *w*_*i*_ are adjusted by gradient descent learning using the development partition. This is done after the training of the base models, using their predictions as input to the above linear model.

#### Learned weights by family (LWF) voting

As a further step along, weights of each model can be learned for each family *k*. In this schema, the weight *w*_*ik*_ is also learned with gradient descent using the development partition. Now, the weighted output of the ensemble is

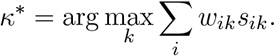

## 3 Data and experimental setup

### 3.1 Data

Pfam data was used for the experiments as in [3], where seed domains of 17,929 families of Pfam v.32.0 were split into train, development (dev) and test sets by clustering based on sequence identity. In order to assure remote homology (low similarity between training and testing sequences), single-linkage clustering at 25% identity within each family was used to build a clustered split. This annotation task provides a realistic scenario close to a real-world annotation problem, where testing domains might not be similar to the training ones. The clustered benchmark was chosen because all the held-out test domains are guaranteed to be far from the training set, avoiding inflated results. However, differently from ProtENN [3], in this work we have retrieved the full-length sequences corresponding to the representative training and testing domains. After that, we eliminated duplicate sequences present both in the training and testing set. The final benchmark dataset has 1,096,510 full protein sequences, with 1,063,492 sequences for training, 16,675 sequences for development and 16,343 sequences for testing.

This full benchmark dataset has a very large class imbalance when each possible Pfam family is considered as the output class. There are several Pfam families with just 1 member sequence, while there are other populated families with thousands of members. For example the family PF13649 appears in more than 4,000 sequences. Then, there is the family PF01660 with just 2 samples, or the family PF02228 with 1 sequence. From a total of 17,929 families, there are just 2,962 families with more than 100 samples for training. There are 8,488 families with training samples between 10 and 100; and 6,478 families with less than 10 training samples each. Figure 2 shows a Venn diagram with the number of families at each partition (train, development and test) of the full dataset. It can be seen that, for example, there are 11 families that are only included in the test partition, that is, those are test families without any sequence to learn from in the training partition. There are 13,285 families with training samples that are not included in development nor test partitions. From a total of 17,929 families, only 2,502 families have samples in train, development and test partition. Nevertheless, these 13,285 families only included in the train partition can be actually an advantage for the DL models, since those families provide a large variety of features and patterns to learn from. This is just opposite to the way HMMs learn, where each model learns only from samples of just one family, independently. Moreover, since there are many extra families in training, each DL model can be specialized in some types of families increasing the diversity of base models, which is well recognized as a property that improves the performance of ensembles [17].

**Figure 2.**
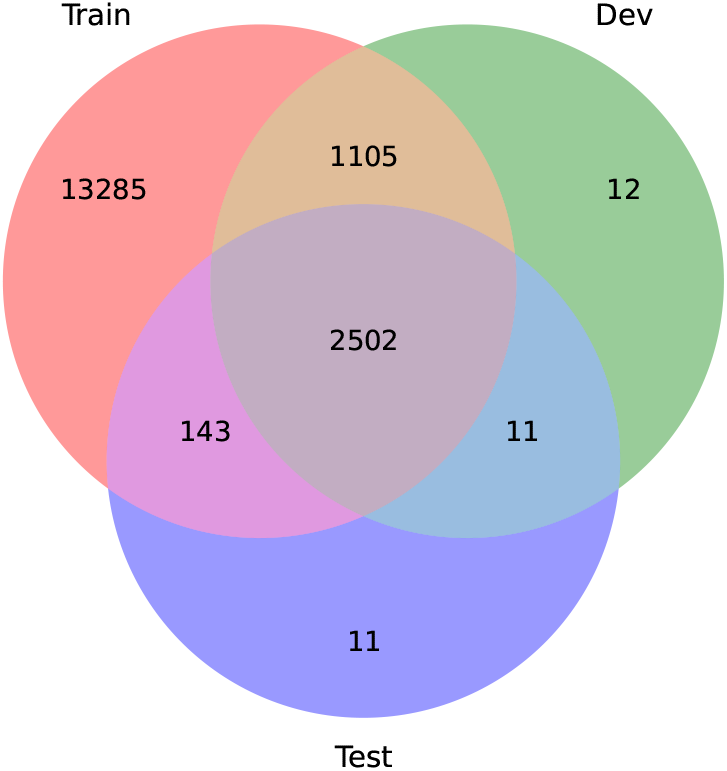
Pfam families class distribution in the full benchmark dataset. The Venn diagram indicates the number of Pfam families in each partition of the dataset: training, development and testing

Due to the very large imbalance in the full dataset partitions, and in order to test the novel proposal first in a more controlled scenario, we have built a subset of the full benchmark dataset and named it “mini dataset”. The mini dataset families distribution was developed in order to guarantee at least 10 sequences for each family in the test partition, which lead to a total of 74,719 proteins and 395 Pfam families that are common to the train, development and test partition. The mini dataset has 56,383 sequences for train, 9,076 sequences for development and 9,260 sequences for test.

### 3.2 Experimental setup

We compared our proposal with the state-of-the-art prediction model currently used at Pfam for family annotation, pHMM [9]. We also compared our proposal with the DL model ProtENN [3], which is based on one-hot encoding inputs and single domains. As a baseline comparison, we included BLASTp [1], one of the most well-known algorithms for similar sequences search.

To ensure a fair comparison with the HMM models, we trained pHMMs from scratch, that is, starting from the multiple sequence alignments (MSAs) required for fitting the HMMs. We created custom MSAs using sequences from the training and development set, aligned with Muscle 3.8.31 [10] using its default settings. Using the MSAs generated, we then employed HMMER 3.4 [9] to build the pHMMs for each family domain and to make the predictions over the test set. It has to be mentioned that in [3] the pHMMs were built using the MSAs already provided by Pfam v32.0 including all the curated seed domains, thus those alignments included seed sequences from the complete dataset (train, dev and test) introducing data leakage in the cross validation.

For obtaining the embeddings we followed the protocol of [22]. The pLM used to obtain the data embeddings was ESM2 [15], in particular esm2 t33 650M UR50D, with size *E* = 1, 280. The window lengths and batch-sizes used in the ensemble members were 32, 64 and 128, with step 4. The development partition was used for early stopping of each DL base model, with patience of 5 epochs. The learning rates explored were *l*_*r*_ *∈* {1E*−*4, 1E*−*5, 1E*−*6}. ResNet was configured with *F* = 1, 100 filters for the first and the last convolutions, and *F*_*b*_ = 550 filters in the bottleneck layer.

For each prediction model or ensemble, the error rate was calculated based on the recall (or sensitivity) as

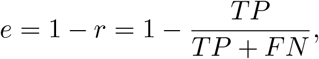

where *TP* are the true positives, and *FN* the false negatives.

## 4 Results

### 4.1 ET-Pfam family prediction in the mini dataset

As an initial evaluation of the ensemble proposal ET-Pfam in a controlled and balanced scenario, we first tested the model and the ensemble strategies in the mini dataset, where all output classes are well represented in the three partitions: train, dev and test. Supplementary Table 1 shows each individual base model hyperparameters, corresponding window length and learning rate, with the errors of each base model for the test partition calculated with the center window (CwS), sliding window area (SwA) and sliding window coverage (SwC) criteria. The best individual model according to CwS reached a 4.72% error, according to SwA the best individual error was 3.64%, and with SwC it was 3.66%.

In order to choose the best ensemble strategy for this dataset, first the errors of each ensemble strategy were calculated on the development partition, with an increasing number of members in the ensemble, from 2 to 10 base models. Figure 3 shows the ET-Pfam ensemble error at the y-axis and the number of base models being incorporated at the x-axis. The top plot shows the different ensemble strategies error when CwS is considered for the base model scores. In this case the best configuration is an ensemble of 10 base models with the LWF voting strategy (error 0.57%). The middle plot shows the ET-Pfam ensemble error when SwA is considered for the base model scores, being here again LWF voting strategy the best one with 10 base models (error 1.54%). Finally the bottom plot presents the ET-Pfam error when SwC is used, achieving best performance (error 1.53%) after 10 models ensembled with LWF, again. In summary, the novel LWS ensemble strategy has reached, in all cases, the best performance. In the best case, a very impressive low error of 0.57% was obtained in this dataset (full details in Supplementary Table 2).

**Figure 3.**
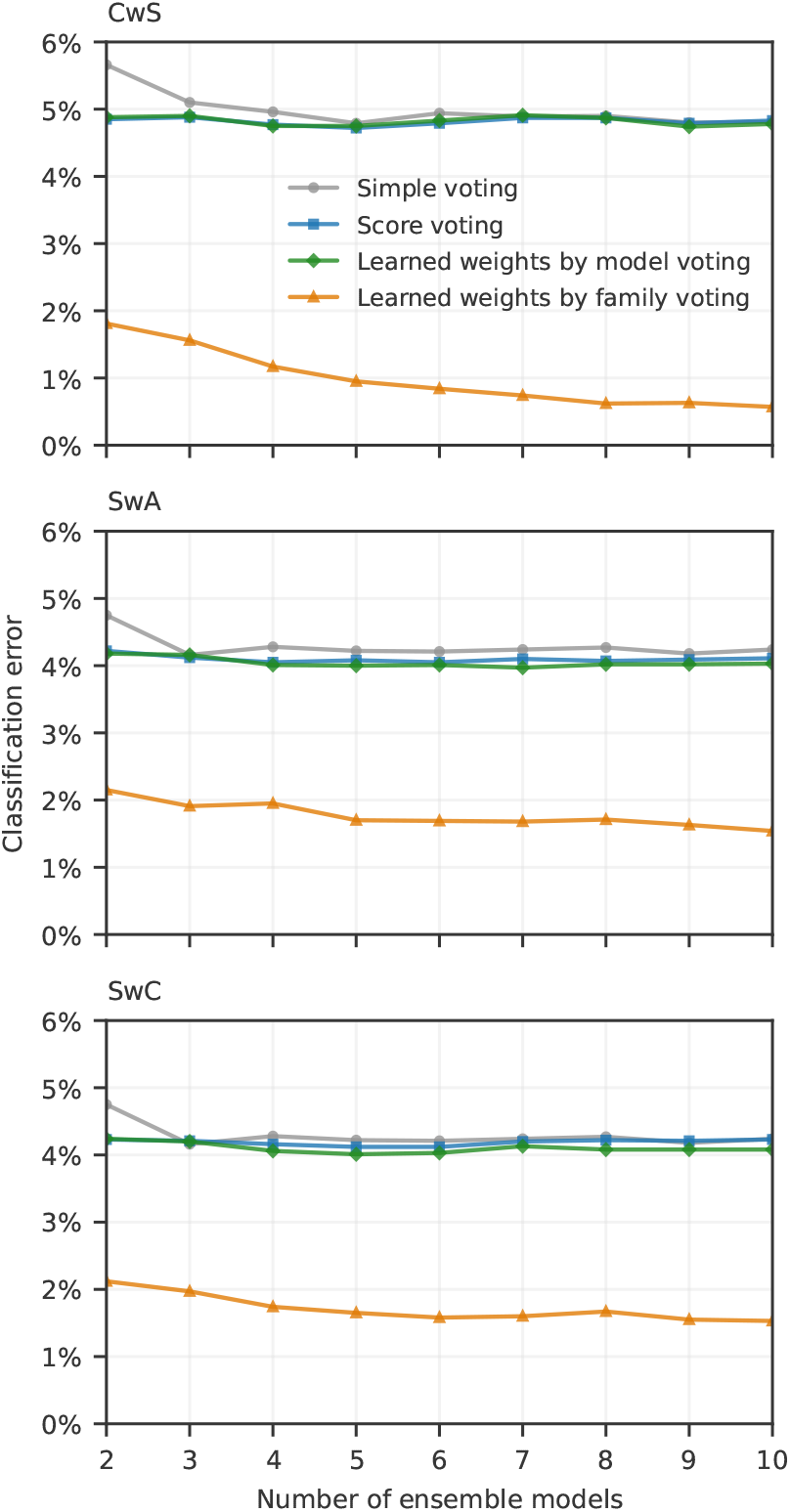
Details of the ET-Pfam classification error for different ensemble strategies (in different colors) at the mini dataset development partition. CwS: centered window score (top); SwA: sliding window area (middle); SwC: sliding window coverage (bottom)

Then, after the best ensemble strategy and number of base models has been chosen using the independent development subset, the base models are ensembled with the winning strategy, ET-Pfam LWF, and tested with the test partition. Table 1 shows that the pHMM error in the mini dataset is 22.70%. Here, the ProtENN+pLM [3] error is 4.40% for CwS, 3.79% for SwA and 3.79% for SwC. When the CwS is considered for the base models error calculation, the ET-Pfam LWF classification error is 1.24%. For the SwA the ET-Pfam LWF error is 1.49%. Finally, for the SwC the ET-Pfam LWF classification error is 1.57%.

**Table 1:**
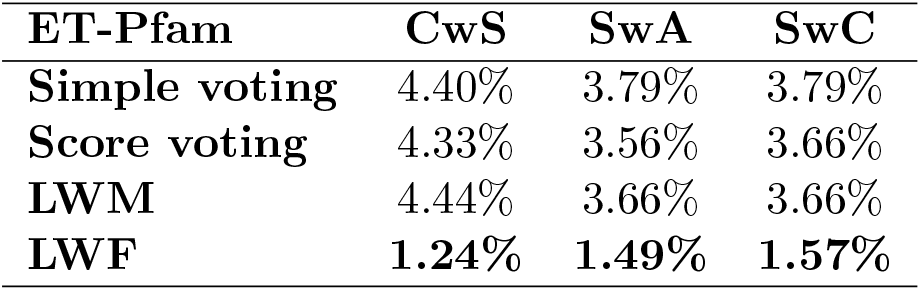
Ensemble strategies error and pHMM error at the test partition for the mini dataset.

### 4.2 ET-Pfam family prediction in the full dataset

Following the same methodology proposed in the previous subsection, we first evaluated each DL base model on the full train dataset, using the dev partition during training for early stopping. Supplementary Table 3 shows in rows each base model to be ensembled, the corresponding window length and learning rate explored. The best individual model in the full dataset (*W* = 128 and *l*_*r*_ = 1.00*E −* 04) achieved a 13.94% error with CwS, a 12.91% error with SwA, and 12.98% with SwC.

In order to choose the best strategy for the full dataset, the errors of each ET-Pfam ensemble strategy were calculated at the development partition while base models were incrementally incorporated into the ensemble, from 2 to 10 (only those models with a central window error below 20%). Table 2 shows the ensemble error according to the number of base models being incorporated at the ensemble, and for all the ensemble strategies. In all cases the best ensemble strategy was using 10 base models and learned weights by family voting strategy (just marginal improvements were obtained from 6 integrated models). The best ET-Pfam ensemble error with CwS (10 models) is achieved with ET-Pfam LWF 0.84%. The ET-Pfam ensemble error with SwA converges to 3.82% with ET-Pfam LWF. Finally, the bottom subtable indicates ensemble error when SwC and ET-Pfam LWF are used, achieving an error of 3.32% with 10 ensembled models.

**Table 2:**
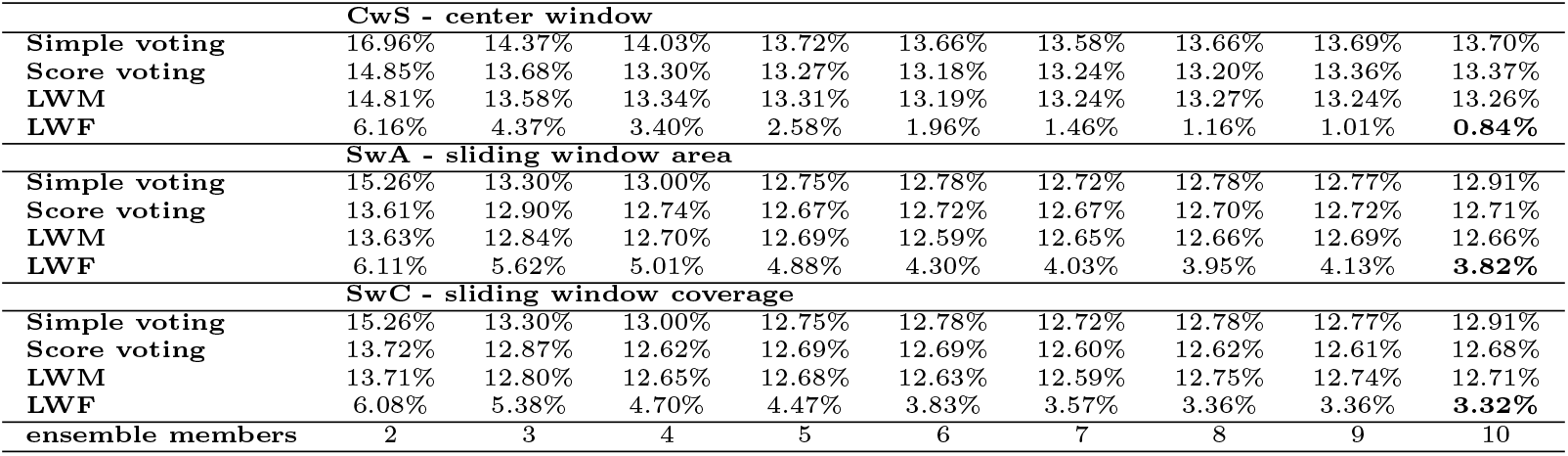
ET-Pfam ensemble strategies error at the development partition for the full dataset as each base model is being incorporated into the ensemble, from 2 to 10 models.

Once the best number of base models (10) has been selected according to the results on the development partition, the DL base models were ensembled and tested in the test partition. Results are provided in Table 3. The ProtENNBileschi2022 error is 18.05%. The pHMM [9] error for the full dataset test partition is 29.28%, while the BLASTp [1] error is 34.94%. It is very interesting to notice that all the ET-Pfam ensemble strategies provide slower errors than the pHMM models, ProtENN and BLASTp, being LWF the ensemble strategy with the lowest error. When the CwS is considered for the base models error calculation, the ET-Pfam LWF classification error is 7.61%. For the SwA, the error is 7.32%, and for the SwC the error is 7.00%, the lowest error, which represents just 1,490 sequences within the complete test set of 21,293 proteins. It has to be highlighted that this error is considerably lower than the best individual base model error of 12.91%. Differently from the mini dataset, where the best error criteria was the same for the development and for the test partition, in the full dataset the best error criteria is the SwC, which is more robust and has a better generalization capacity in this much larger dataset.

**Table 3:**
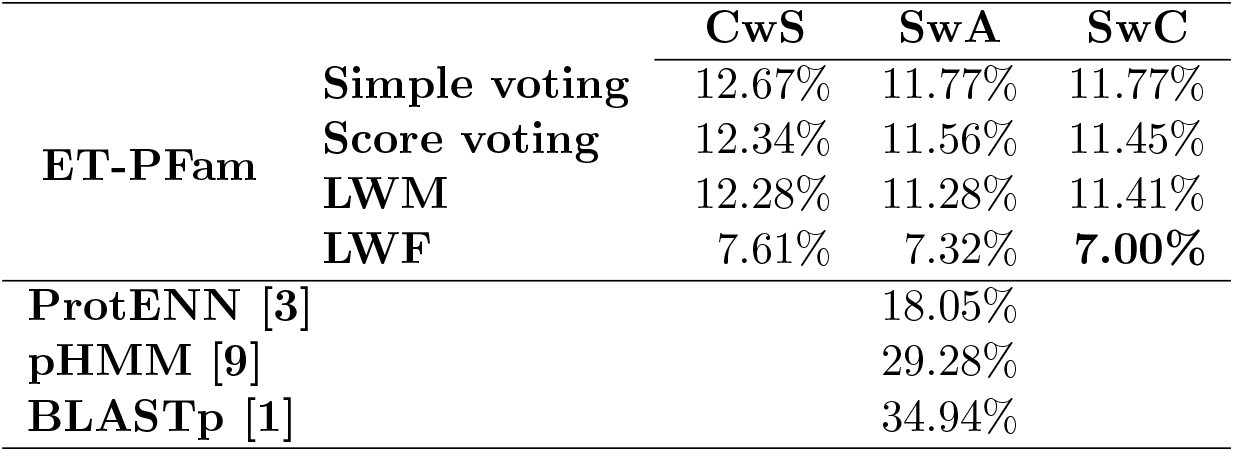
ET-Pfam ensemble strategy and competitor errors at the test partition for the full dataset. Best performance (lowest error) in bold.

|  |  | **CwS** | **SwA** | **SwC** |
| --- | --- | --- | --- | --- |
| **ET-Pfam** | **Simple voting** | 12.67% | 11.77% | 11.77% |
|  | **Score voting** | 12.34% | 11.56% | 11.45% |
|  | **LWM** | 12.28% | 11.28% | 11.41% |
|  | **LWF** | 7.61% | 7.32% | **7.00%** |
| **ProtENN [3]** |  |  | 18.05% |  |
| **pHMM [9]** |  |  | 29.28% |  |
| **BLASTp [1]** |  |  | 34.94% |  |

In summary, comparative results of our proposal ET-Pfam and the state-of-the-art procedure currently used at Pfam for protein annotations has shown that ET-Pfam can provide effectively improved results, reducing the prediction error by an impressive four times in comparison to the pHMM and BLASTp techniques, and by more than half in comparison to the DL competitor. Our proposal is an ensemble of DL models together with transfer learning from pLM embeddings and a novel ensemble strategy that leverages the per-family models predictions, significantly diminishing the prediction error when new proteins have to be annotated with the Pfam family domains.

## 5 Conclusions

This work has presented ET-Pfam, a novel approach for Pfam family prediction based on an ensemble of diverse DL base models that use transfer learning. A novel ensemble strategy learns weights for voting with the per-family output scores of DL models. Comparative results versus the classical HMM models used by Pfam and individual DL models highlighted the effectiveness of the proposed ensemble method in improving protein sequence classification, reducing the prediction error four times. Incorporating family-specific weights at the ensemble allowed the base models to better contribute to the Pfam classification. This could not have been possible with a single base model, or with the simple and classical voting combination of base models.

Results achieved have demonstrated the practical utility of embeddings from protein large language models, together with ensembles of deep learning models, for the Pfam annotation problem. We presented here a substantial advance over previous efforts applying deep learning in terms of improving the state of the art prediction and the rigorous comparison with existing methods. The next step is to use the ET-Pfam model presented here to rapidly and efficiently annotate unlabeled sequences with the aim of further expanding the annotated protein universe, which could be of special interest for biotechnology and health applications. Future work will also focus on investigating the output scores for automatic segmentation of domains.

## Supporting information

Supplementary Maerial

## 7 Author contributions statement

G.S., L.A.B. and D.H.M conceived the research goals, designed the models and developed the experimental methodology; S.E., S.A.D., R.V. and E.F. wrote the code and conducted the experiments; S.A.D. prepared the figures; G.S. wrote the original draft; G.S. and D.H.M. analysed the results and wrote the manuscript; all authors reviewed, edited and approved the manuscript.

## References

[1] S. Altschul. Gapped blast and psi-blast: a new generation of protein database search programs. Nucleic Acids Research, 25(17):3389–3402, September 1997.

[2] A Bateman and et al. The pfam protein families database. Nucleic Acids Research, 32(90001):138D – 141, 2004.

[3] M Bileschi and et al. Using deep learning to annotate the protein universe. Nature Biotechnology, 40(6):932–937, 2022.

[4] L Bugnon and et al. Transfer learning: The key to functionally annotate the protein universe. Patterns, 4(2):100691, 2023.

[5] Y Cao and et al. Ensemble deep learning in bioinformatics. Nature Machine Intelligence, 2(9):500–508, 2020.

[6] N Detlefsen, S Hauberg, and W Boomsma. Learning meaningful representations of protein sequences. Nature Communications, 13(1):1–9, 2022.

[7] A Edera E Fenoy and G Stegmayer. Transfer learning in proteins: evaluating novel protein learned representations for bioinformatics tasks. Briefings in Bioinformatics, 23(4):bbac232, 2022.

[8] S Eddy. Profile hidden markov models. Bioinformatics, 14(9):755–763, 1998.

[9] S Eddy. Accelerated profile hmm searches. PLOS Computational Biology, 7(10):1–16, 10 2011.

[10] R Edgar. Muscle: a multiple sequence alignment method with reduced time and space complexity. BMC Bioinformatics, 5(1):1–10, 2004.

[11] M Ganaie and et al. Ensemble deep learning: A review. Engineering Applications of Artificial Intelligence, 115:105151, 2022.

[12] M Heinzinger and B Rost. Teaching ai to speak protein. Current Opinion in Structural Biology, 91:102986, 2025.

[13] T Idhaya, A Suruliandi, and S Raja. A comprehensive review on machine learning techniques for protein family prediction. The Protein Journal, 43(2):171–186, 2024.

[14] J Jumper and et al. Highly accurate protein structure prediction with alphafold. Nature, 596(7873):583–589, 2021.

[15] Z Lin and et al. Evolutionary-scale prediction of atomic-level protein structure with a language model. Science, 379(6637):1123–1130, 2023.

[16] T Paysan-Lafosse and et al. The pfam protein families database: embracing ai/ml. Nucleic Acids Research, 53(D1):D523–D534, 2024.

[17] M Pividori, G Stegmayer, and D Milone. Diversity control for improving the analysis of consensus clustering. Information Sciences, 361-362:120– 134, 2016.

[18] A Rives and et al. Biological structure and function emerge from scaling unsupervised learning to 250 million protein sequences. Proceedings of the National Academy of Sciences, 118(15):e2016239118, 2021.

[19] S Seo and et al. Deepfam: deep learning based alignment-free method for protein family modeling and prediction. Bioinformatics, 34(13):i254–i262, 2018.

[20] C Tran, S Khadkikar, and A Porollo. Survey of protein sequence embedding models. International Journal of Molecular Sciences, 24(4):3775, 2023.

[21] S Unsal and et al. Learning functional properties of proteins with language models. Nature Machine Intelligence, 4(3):227–245, 2022.

[22] R Vitale and et al. Evaluating large language models for annotating proteins. Briefings in Bioinformatics, 25(3):bbae177, 2024.

[23] L Wand and T Jiang. On the complexity of multiple sequence alignment. Journal of Computational Biology, 1(4):337–348, 1994.

[24] K Weiss, T Khoshgoftaar, and D Wang. A survey of transfer learning. Journal of Big Data, 3(1):1–11, 2016.

[25] K Weissenow and B Rost. Are protein language models the new universal key? Current Opinion in Structural Biology, 91:102997, 2025.

[26] K Yang and et al. Learned protein embeddings for machine learning. Bioinformatics, 34(15):2642–2648, 2018.

