## Supplementary Maerial for "ET-Pfam: Ensemble transfer learning for protein family prediction"

### 1 Pfam family prediction in the mini dataset

Table S1: Individual models error according to several possible base model score calculation criteria at the test partition for the mini dataset. Best model with lowest error in bold.

| **Model** | $W$ | $l_r$ | **CwS** | **SwA** | **SwC** |
|---|---|---|---|---|---|
| 1 | 32 | 1.00E-04 | 5.66% | 3.73% | 3.92% |
| 2 | | 1.00E-05 | 5.51% | 3.90% | 4.02% |
| 3 | | 1.00E-06 | 6.13% | 4.12% | 4.33% |
| 4 | | | 5.48% | **3.64%** | **3.66%** |
| 5 | | | 5.68% | 3.99% | 3.98% |
| 6 | | | 5.65% | 3.85% | 3.90% |
| 7 | 64 | 1.00E-06 | 5.27% | 3.65% | 3.74% |
| 8 | | | 5.15% | 3.88% | 3.87% |
| 9 | 128 | 1.00E-06 | **4.72%** | 4.22% | 4.40% |
| 10 | | | 4.88% | 4.05% | 4.21% |

Table S2: Ensemble strategies error at the development partition for the mini dataset as each base model is being incorporated into the ensemble, from 2 to 10 models.

| | **2** | **3** | **4** | **5** | **6** | **7** | **8** | **9** | **10** |
|---|---|---|---|---|---|---|---|---|---|
| **CwS - center window** | | | | | | | | | |
| **Simple voting** | 5.66% | 5.10% | 4.96% | 4.79% | 4.94% | 4.89% | 4.90% | 4.80% | 4.81% |
| **Score voting** | 4.85% | 4.88% | 4.77% | 4.72% | 4.79% | 4.87% | 4.87% | 4-79% | 4.83% |
| **LWM** | 4.88% | 4.90% | 4.75% | 4.75% | 4.83% | 4.91% | 4.87% | 4.74% | 4.78% |
| **LWF** | 1.81% | 1.56% | 1.17% | 0.95% | 0.84% | 0.74% | 0.62% | 0.63% | **0.57%** |
| **SwA - sliding window area** | | | | | | | | | |
| **Simple voting** | 4.75% | 4.16% | 4.28% | 4.22% | 4.21% | 4.24% | 4.27% | 4.18% | 4.24% |
| **Score voting** | 4.22% | 4.12% | 4.05% | 4.08% | 4.05% | 4.10% | 4.07% | 4.09% | 4.11% |
| **LWM** | 4.18% | 4.16% | 4.01% | 4.00% | 4.01% | 3.97% | 4.02% | 4.02% | 4.03% |
| **LWF** | 2.15% | 1.91% | 1.95% | 1.70% | 1.69% | 1.68% | 1.71% | 1.63% | **1.54%** |
| **SwC - sliding window coverage** | | | | | | | | | |
| **Simple voting** | 4.75% | 4.16% | 4.28% | 4.22% | 4.21% | 4.24% | 4.27% | 4.18% | 4.24% |
| **Score voting** | 4.23% | 4.21% | 4.16% | 4.12% | 4.12% | 4.20% | 4.22% | 4.21% | 4.23% |
| **LWM** | 4.24% | 4.20% | 4.06% | 4.01% | 4.03% | 4.13% | 4.08% | 4.08% | 4.08% |
| **LWF** | 2.12% | 1.97% | 1.74% | 1.65% | 1.58% | 1.60% | 1.67% | 1.55% | **1.53%** |

Table S3: Ensemble strategies error and pHMM error at the test partition for the mini dataset.

| **ET-Pfam ensemble strategy** | **CwS** | **SwA** | **SwC** |
|---|---|---|---|
| **Simple voting** | 4.40% | 3.79% | 3.79% |
| **Score voting** | 4.33% | 3.56% | 3.66% |
| **LWM** | 4.44% | 3.66% | 3.66% |
| **LWF** | **1.24%** | **1.49%** | **1.57%** |
| **pHMM** | | 22.70% | |

### 2 Pfam family prediction in the full dataset

Table S4: Individual models error according to several possible base model score calculation criteria at the test partition for the full dataset. Best model, with lowest error, in bold.

| **Model** | $W$ | $l_r$ | **CwS** | **SwA** | **SwC** |
|---|---|---|---|---|---|
| 1 | | 1.00E-04 | 25.87% | 20.86% | 21.34% |
| 2 | 32 | 1.00E-05 | 22.76% | 18.18% | 18.49% |
| 3 | | 1.00E-06 | 27.00% | 20.56% | 21.18% |
| 4 | | 1.00E-04 | 21.95% | 18.67% | 18.81% |
| 5 | | 2.00E-04 | 18.94% | 15.77% | 16.02% |
| 6 | | 3.00E-04 | 26.64% | 21.30% | 21.72% |
| 7 | 64 | 1.00E-05 | 20.94% | 17.00% | 17.42% |
| 8 | | 2.00E-05 | 21.59% | 18.26% | 18.45% |
| 9 | | 3.00E-05 | 21.56% | 18.29% | 18.11% |
| 10 | | 1.00E-06 | 21.23% | 16.74% | 17.24% |
| 11 | | 1.00E-04 | 18.46% | 16.17% | 16.31% |
| 12 | | 2.00E-04 | 21.05% | 18.26% | 18.55% |
| 13 | | 3.00E-04 | 16.17% | 14.20% | 14.22% |
| 14 | | 3.00E-04 | 15.74% | 13.75% | 13.91% |
| 15 | | 1.00E-04 | **13.94%** | **12.91%** | **12.98%** |
| 16 | | 1.00E-04 | 14.73% | 13.30% | 13.07% |
| 17 | | 1.00E-04 | 15.00% | 13.03% | 12.99% |
| 18 | | 1.00E-04 | 14.72% | 13.56% | 13.43% |
| 19 | 128 | 1.00E-05 | 15.44% | 14.04% | 14.28% |
| 20 | | 1.00E-05 | 15.36% | 13.88% | 13.74% |
| 21 | | 1.00E-05 | 15.64% | 13.97% | 14.08% |
| 22 | | 1.00E-05 | 14.94% | 13.69% | 13.58% |
| 23 | | 1.00E-05 | 16.79% | 14.77% | 15.11% |
| 24 | | 1.00E-05 | 16.94% | 15.24% | 15.50% |
| 25 | | 2.00E-05 | 16.89% | 14.97% | 15.56% |
| 26 | | 3.00E-05 | 17.30% | 15.64% | 15.89% |
| 27 | | 1.00E-06 | 20.31% | 16.84% | 17.45% |

Table S5: ET-Pfam ensemble strategies error at the development partition for the full dataset as each base model is being incorporated into the ensemble, from 2 to 10 models.

| | **2** | **3** | **4** | **5** | **6** | **7** | **8** | **9** | **10** |
|---|---|---|---|---|---|---|---|---|---|
| **CwS - center window** | | | | | | | | | |
| **Simple voting** | 16.96% | 14.37% | 14.03% | 13.72% | 13.66% | 13.58% | 13.66% | 13.69% | 13.70% |
| **Score voting** | 14.85% | 13.68% | 13.30% | 13.27% | 13.18% | 13.24% | 13.20% | 13.36% | 13.37% |
| **LWM** | 14.81% | 13.58% | 13.34% | 13.31% | 13.19% | 13.24% | 13.27% | 13.24% | 13.26% |
| **LWF** | 6.16% | 4.37% | 3.40% | 2.58% | 1.96% | 1.46% | 1.16% | 1.01% | **0.84%** |
| **SwA - sliding window area** | | | | | | | | | |
| **Simple voting** | 15.26% | 13.30% | 13.00% | 12.75% | 12.78% | 12.72% | 12.78% | 12.77% | 12.91% |
| **Score voting** | 13.61% | 12.90% | 12.74% | 12.67% | 12.72% | 12.67% | 12.70% | 12.72% | 12.71% |
| **LWM** | 13.63% | 12.84% | 12.70% | 12.69% | 12.59% | 12.65% | 12.66% | 12.69% | 12.66% |
| **LWF** | 6.11% | 5.62% | 5.01% | 4.88% | 4.30% | 4.03% | 3.95% | 4.13% | **3.82%** |
| **SwC - sliding window coverage** | | | | | | | | | |
| **Simple voting** | 15.26% | 13.30% | 13.00% | 12.75% | 12.78% | 12.72% | 12.78% | 12.77% | 12.91% |
| **Score voting** | 13.72% | 12.87% | 12.62% | 12.69% | 12.69% | 12.60% | 12.62% | 12.61% | 12.68% |
| **LWM** | 13.71% | 12.80% | 12.65% | 12.68% | 12.63% | 12.59% | 12.75% | 12.74% | 12.71% |
| **LWF** | 6.08% | 5.38% | 4.70% | 4.47% | 3.83% | 3.57% | 3.36% | 3.36% | **3.32%** |
